# Increased infection risk in Addison’s disease and congenital adrenal hyperplasia: a primary care database cohort study

**DOI:** 10.1101/628156

**Authors:** Alberto S. Tresoldi, Dana Sumilo, Mary Perrins, Konstantinos A. Toulis, Alessandro Prete, Narendra Reddy, John A.H. Wass, Wiebke Arlt, Krishnarajah Nirantharakumar

## Abstract

**Context:** Mortality and infection-related hospital admissions are increased in patients with primary adrenal insufficiency (PAI). However, the risk of primary care-managed infections in patients with PAI is unknown.

**Objective:** To estimate infection risk in PAI due to Addison’s disease (AD) and congenital adrenal hyperplasia (CAH) in a primary care setting.

**Design:** Retrospective cohort study using UK data collected from 1995 to 2018.

**Main outcome measures:** Incidence of lower respiratory tract infections (LRTIs), urinary tract infections (UTIs), gastrointestinal infections (GIIs), and prescription counts of antimicrobials in adult PAI patients compared to unexposed controls.

**Results:** A diagnosis of PAI was established in 1580 AD patients (mean age 51.7 years) and 602 CAH patients (mean age 35.4 years). All AD patients and 42% of CAH patients were prescribed glucocorticoids, most frequently hydrocortisone in AD (82%) and prednisolone in CAH (50%). AD and CAH patients exposed to glucocorticoids, but not CAH patients without glucocorticoid treatment, had a significantly increased risk of LRTIs (adjusted incidence rate ratio AD 2.11 [95% confidence interval 1.64-2.69], CAH 3.23 [1.21-8.61]), UTIs (AD 1.51 [1.29-1.77], CAH 2.20 [1.43-3.34]), and GIIs (AD 3.80 [2.99-4.84], CAH 1.93 [1.06-3.52]). This was mirrored by increased prescription of antibiotics (AD 1.73 [1.69-1.77], CAH 1.77 [1.66-1.89]) and antifungals (AD 1.89 [1.74-2.05], CAH 1.91 [1.50-2.43]).

**Conclusions:** There is an increased risk of infections and antimicrobial use in PAI in the primary care setting at least partially linked to glucocorticoid treatment. Future studies will need to address whether more physiological glucocorticoid replacement modes could reduce this risk.

**Précis:** Using data from 1580 AD patients and 602 CAH patients collected in a UK primary care database from 1995 to 2018, we identified increased risk of infections and antimicrobial prescription counts.

## INTRODUCTION

Primary adrenal insufficiency (PAI) is a severe and potentially life-threatening condition caused by the failure of the adrenal cortex to produce glucocorticoids and, in most cases, mineralocorticoids, which occurs in the setting of adrenal disease (1). The two most frequent causes of PAI are autoimmune adrenalitis, the most frequent cause of Addison’s disease, AD, in Western countries, and congenital adrenal hyperplasia (CAH).

The prognosis of patients with PAI has improved considerably after life-saving glucocorticoid replacement therapy became available in the 1950s; however, an increased risk of death has been described in both AD and CAH patients even in recent years (2,3). In patients with AD, this has been attributed to adrenal crisis- and infection-related mortality (4), while for both CAH and AD patients an increased cardiovascular-related mortality has been described (3,4). Other studies have reported an increased use of antimicrobial agents and infection-related hospital admissions in patients with PAI (5,6). Recent evidence suggests that the increased risk of infections in these patients could be explained by an impairment of natural killer cell function (7), which may be caused by the non-physiological delivery of glucocorticoids by currently available preparations and an associated change in clock gene expression patterns in immune cells (7,8). No studies have estimated yet the overall risk of common infections in people with PAI, i.e. infections that are primarily managed in the primary care setting and usually do not require hospital admission. However, such infections potentially expose PAI patients to significant risk of adrenal crisis. Therefore, this study aimed to assess the risk of common types of primary care-managed infections, namely infections of the lower respiratory tract, the urinary tract, and gastrointestinal infections, and the use of antimicrobials in the primary care setting in patients with PAI, including both AD and CAH patients with and without glucocorticoid therapy.

## MATERIALS AND METHODS

### Study design and setting

We conducted a population-based, retrospective, open cohort study to determine the infection risk of patients with AD and CAH in the primary care setting. We assessed the risk of lower respiratory tract infections (LRTIs), urinary tract infections (UTIs), gastrointestinal infections (GIIs), and the counts of antimicrobial prescriptions. We used data from The Health Improvement Network (THIN) database, comprising anonymized electronic medical records from UK general practitioner (GP) practices covering over 5% of the UK population. THIN holds data on demographic characteristics, clinical diagnoses, physical measurements, laboratory results and drug prescriptions recorded using clinical Read code system. Patients registered in THIN have similar age and sex distributions to the general UK population and, therefore, THIN data are well suited for epidemiological studies (9,10).

### Study population and period

Our study population consisted of two “exposed” cohorts, comprising adult patients (≥18 years old) diagnosed with AD or CAH according to selected Read codes (see Appendix for the codes used) (11,12). We excluded patients who were at any time point coded with a code consistent with other causes of adrenal insufficiency. We could not retrieve data on 21-hydroxylase autoantibodies in the study participants, due to the nature of the study. Therefore, in this paper, we defined AD as primary adrenal insufficiency not caused by CAH. To ensure accuracy of case definition in the AD cohort we only included patients who had at least one prescription of both glucocorticoids (accepting glucocorticoids commonly used in AD) and mineralocorticoids. We also performed a sensitivity analysis to include only patients who had at least two prescription of both glucocorticoids and mineralocorticoids. We subdivided the CAH cohort in two sub-cohorts: (a) patients who had at least one glucocorticoid prescription at any point (using the same glucocorticoid codes used for AD patients) and (b) patients who were never prescribed with any glucocorticoid therapy, since patients with CAH do not always require glucocorticoid therapy. For every exposed patient, we randomly selected two individuals from a pool of patients matched for age, sex and GP practice who did not have a Read code consistent with PAI at any point before or during the observation period.

The study period extended from 1 January 1995 to 1 January 2018. Patients were eligible for inclusion one year from the latest of the following dates: study start date, patient registration date with the GP practice, and practice eligibility date (the date when practices have implemented an electronic medical record and have passed the assessment for acceptable data quality). The one-year lag period was applied to ensure there was enough time to document all information accurately after registration with the practice or after a practice was deemed eligible to take part. To ensure acceptable data quality, practices were required to have used the electronical health record system for at least one year and have acceptable mortality reporting (13).

The index date, i.e. the date when follow-up commences, was defined as the date of diagnosis for newly diagnosed patients or, if they were already diagnosed with PAI, the date when they registered with an eligible GP practice. Patients were followed from index date up until the earliest of the following dates: outcome of interest (only for estimating the incidence of infections), patient transfer date from practice, patient death, practice’s last data collection date, and study end date.

### Outcomes

For the first outcome, the incidence of infections, we used the Read codes that identify cases of LRTIs, UTIs and GIIs (see Suppl. Appendix). These infections were chosen because they are the most common type of infections evidenced in general population and they are frequently diagnosed in primary care (14). We then calculated the occurrence of this outcome in the different cohorts.

For the second outcome, antimicrobial use, we used the codes for antibiotics and antifungals as classified in the British National Formulary. We then calculated the total number of prescriptions for every antimicrobial in each cohort.

For each of the study groups we analyzed age, sex, body mass index (BMI), smoking status, Townsend Deprivation Index (a measure of deprivation within a population) (15), Charlson Comorbidity Index (a method of classifying comorbidities to predict mortality in primary care) (16,17), and type of glucocorticoids prescribed at baseline. For the AD cohort (in which most patients were likely to have autoimmune PAI), given the frequent association with other autoimmune conditions, we also evaluated the prevalence of associated autoimmune comorbidities.

### Statistical analysis

Descriptive statistics were used to summarize the baseline characteristics for the exposed and unexposed groups of patients. Categorical variables were investigated using Chi-square test and continuous variables were analyzed using t-test.

Adjusted incidence rate ratios (aIRRs) for specific infections and antimicrobial prescriptions were calculated after adjustment for age, sex, smoking status, BMI, Townsend Deprivation Index, and Charlson Comorbidity Index, using multivariate Poisson regression analysis. Statistical analyses were conducted using Stata version 14.2 (Stata Corp, College Station Texas, USA) and GraphPad Prism 7.04 (GraphPad Software Inc, San Diego, CA).

### Ethical approval

The THIN database obtained ethical approval from the South East Multicentre Research Ethics Committee in 2003. The present study was reviewed and approved (study reference: 18THIN063) by the THIN Scientific Review Committee in July 2018.

## RESULTS

### Baseline characteristics of the AD cohort

In total, 1580 patients fulfilled the AD criteria; these were matched with 3158 unexposed individuals (Table 1). The mean age of AD patients was 51.7 years and the majority were women (57.8%). Compared to unexposed individuals, AD patients had a lower median BMI, while the Townsend Deprivation Index did not differ significantly between the two groups. The Charlson Comorbidity Index showed that AD patients had an increased burden of comorbidities compared to the matched population; this included a higher prevalence of autoimmune comorbidities, including autoimmune thyroid diseases, type 1 diabetes mellitus, ulcerative colitis, celiac disease and pernicious anemia (Table 1).

**Table 1:** Baseline characteristics of patients with Addison’s disease (AD) and matched unexposed patients.

|  | **AD patients**<br>**(n = 1580)** | **Matched unexposed**<br>**patients (n = 3158)** |
| --- | --- | --- |
| **Age** years, mean ± SD | 51.7 ± 18.5 | 51.7 ± 18.5 |
| **Male sex**, n. (%) | 666 (42.2%) | 1.330 (42.1%) |
| **BMI** | <u>Total n = 1227</u> | <u>Total n = 2264</u> |
| median (IQR) | 24.3 (21.5-27.6)* | 25.5 (22.6-28.9) |
| □ 18.5 kg/m<sup>2</sup>, n. (%) | 87 (7.1%) | 51 (2.3%) |
| 18.5-25 kg/m<sup>2</sup>, n. (%) | 606 (49.4%) | 973 (43.0%) |
| 25-30 kg/m<sup>2</sup>, n. (%) | 362 (29.5%) | 791 (34.9%) |
| ≥30 kg/m<sup>2</sup>, n. (%) | 172 (14.0%) | 449 (19.8%) |
| Missing, n. | 353 | 894 |
| **Smoking status** | <u>Total = 1384</u> | <u>Total = 2679</u> |
| Non-smoker, n. (%) | 882 (63.7%)* | 1548 (57.8%) |
| Ex-smoker, n. (%) | 250 (18.1%) | 490 (18.3%) |
| Smoker, n. (%) | 252 (18.2%)* | 641 (23.9%) |
| Missing, n. | 196 | 479 |
| **Townsend Deprivation Index** | <u>Total = 1373</u> | <u>Total = 2780</u> |
| 1 (least deprived), n. (%) | 350 (25.5%) | 743 (26.7%) |
| 2, n. (%) | 306 (22.3%) | 590 (21.2%) |
| 3, n. (%) | 290 (21.1%) | 607 (21.8%) |
| 4, n. (%) | 255 (18.6%) | 471 (16.9%) |
| 5 (most deprived), n. (%) | 172 (12.5%) | 369 (13.3%) |
| Missing, n. | 207 | 378 |
| **Charlson Comorbidity Index** |  |  |
| 0 (no comorbidities), n. (%) | 863 (54.6%)* | 2263 (71.7%) |
| 1, n. (%) | 377 (23.9%)* | 536 (17.0%) |
| ≥2 (more comorbidities), n. (%) | 340 (21.5%)* | 359 (11.4%) |
| **Associated autoimmune comorbidities** |  |  |
| Hyperthyroidism, n. (%) | 40 (2.5%)* | 16 (0.5%) |
| Hypothyroidism, n. (%) | 457 (28.9%)* | 122 (3.9%) |
| Rheumatoid arthritis, n. (%) | 25 (1.6%) | 37 (1.2%) |
| Type 1 diabetes mellitus, n. (%) | 127 (8.0%)* | 15 (0.5%) |
| Inflammatory bowel disease, n. (%) | 29 (1.8%)* | 22 (0.7%) |
| Crohn's disease, n. (%) | 9 (0.6%) | 9 (0.3%) |
| Ulcerative colitis, n. (%) | 20 (1.3%)* | 11 (0.4%) |
| Coeliac disease, n. (%) | 25 (1.6%)* | 10 (0.3%) |
| Multiple sclerosis, n. (%) | n < 5 | 8 (0.3%) |
| Pernicious anaemia, n. (%) | 41 (2.6%)* | 15 (0.5%) |
| Systemic lupus erythematosus, n. (%) | 6 (0.4%) | n < 5 |

\*p <0.05 for AD patients vs. unexposed cohort.

### Baseline characteristics of the CAH cohort

In total, 602 patients fulfilled the CAH criteria and were subdivided into 254 glucocorticoid-treated patients (42.2%) and 348 patients not on glucocorticoids (57.8%). These were matched with 508 and 696 unexposed controls, respectively (Table 2).

**Table 2:** Baseline characteristics of the CAH patients and matched unexposed patients.

|  | CAH cohort<br>(n = 602) | Matched<br>unexposed cohort<br>(n = 1204) | CAH cohort on<br>glucocorticoids<br>(n = 254) | Matched<br>unexposed<br>cohort (n = 508) | CAH cohort not on<br>glucocorticoids<br>(n = 348) | Matched<br>unexposed<br>cohort (n = 69) |
| --- | --- | --- | --- | --- | --- | --- |
| **Age**, years, mean ± SD | 35.4 ± 16.3 | 35.5 ± 16.2 | 33.4 ± 15.1 | 33.5 ± 15.0 | 36.9 ± 17.0<sup>†</sup> | 37.0 ± 16.9 |
| **Male sex**, n. (%) | 167 (27.7%) | 334 (27.7%) | 80 (31.5%) | 160 (31.5%) | 87 (25.0%) | 174 (25.0%) |
| **BMI** | <u>Total = 438</u> | <u>Total = 835</u> | <u>Total = 186</u> | <u>Total = 340</u> | <u>Total = 252</u> | <u>Total = 495</u> |
| median (IQR) | 26.9 (23.2-31.2)* | 24.0 (21.0-28.0) | 27.0 (23.2-32.0)* | 24.0 (21.3-27.9) | 26.9 (23.2-30.9)* | 24.4 (21.8-28.0) |
| □18.5, n. (%) | 12 (2.7%) | 35 (4.2%) | 8 (4.3%) | 9 (2.6%) | 4 (1.6%) | 26 (5.2%) |
| 18.5-25, n. (%) | 162 (37.0%) | 431 (51.6%) | 68 (36.6%) | 187 (55.0%) | 94 (37.3%) | 244 (49.3%) |
| 25-30, n. (%) | 133 (30.4%) | 224 (26.8%) | 52 (28.0%) | 90 (26.5%) | 81 (32.1%) | 134 (27.1%) |
| ≥30, n. (%) | 131 (29.9%) | 145 (17.4%) | 58 (31.2%) | 54 (15.9%) | 73 (29.0%) | 91 (18.4%) |
| Missing, n. | 164 | 369 | 68 | 168 | 96 | 201 |
| **Smoking status** | <u>Total = 524</u> | <u>Total = 1048</u> | <u>Total = 212</u> | <u>Total = 423</u> | <u>Total = 312</u> | <u>Total = 625</u> |
| Non-smoker, n. (%) | 349 (66.6%) | 670 (63.9%) | 138 (65.1%) | 277 (65.5%) | 211 (67.6%) | 393 (62.9%) |
| Ex-smoker, n. (%) | 71 (13.6%) | 142 (13.6%) | 28 (13.2%) | 50 (11.8%) | 43 (13.8%) | 92 (14.7%) |
| Smoker, n. (%) | 104 (19.9%) | 236 (22.5%) | 46 (21.7%) | 96 (22.7%) | 58 (18.6%) | 140 (22.4%) |
| Missing, n. | 78 | 156 | 42 | 85 | 36 | 71 |
| **Townsend Deprivation Index** | <u>Total = 516</u> | <u>Total = 1057</u> | <u>Total = 223</u> | <u>Total = 459</u> | <u>Total = 293</u> | <u>Total = 598</u> |
| 1 (least deprived), n. (%) | 123 (23.8%) | 235 (22.2%) | 52 (23.3%) | 107 (23.3%) | 71 (24.2%) | 128 (21.4%) |
| 2, n. (%) | 107 (20.7%) | 222 (21.0%) | 46 (20.6%) | 92 (20.0%) | 61 (20.8%) | 130 (21.7%) |
| 3, n. (%) | 114 (22.1%) | 232 (22.0%) | 51 (22.9%) | 99 (21.6%) | 63 (21.5%) | 133 (22.2%) |
| 4, n. (%) | 104 (20.2%) | 225 (21.3%) | 44 (19.7%) | 98 (21.4%) | 60 (20.5%) | 127 (21.2%) |
| 5 (most deprived), n. (%) | 68 (13.2%) | 143 (13.5%) | 30 (13.5%) | 63 (13.7%) | 38 (13.0%) | 80 (13.4%) |
| Missing, n. | 86 | 147 | 31 | 49 | 55 | 98 |
| **Charlson Comorbidity Index** |  |  |  |  |  |  |
| 0 (no comorbidities), n. (%) | 457 (75.9%) | 907 (75.3%) | 187 (73.6%) | 394 (77.6%) | 270 (77.6%) | 513 (73.7%) |
| 1, n. (%) | 103 (17.1%) | 242 (20.1%) | 50 (19.7%) | 96 (18.9%) | 53 (15.2%) | 146 (21.0%) |
| ≥2 (more comorbidities), n. (%) | 42 (7.0%) | 55 (4.6%) | 17 (6.7%) | 18 (3.5%) | 25 (7.2%) | 37 (5.3%) |

\*p <0.05 for CAH patients vs. unexposed cohort.
<sup>†</sup>p <0.05 for CAH patients on glucocorticoids vs. CAH patients not on glucocorticoids.

The majority of CAH patients were female (72.3%), with a lower mean age in glucocorticoid-treated patients at cohort entry (33.4 vs. 36.9 years). CAH patients had a higher median BMI compared to controls, and this was evident for both glucocorticoid-treated sub-cohort and the CAH sub-cohort never treated with glucocorticoids. CAH patients were more frequently overweight or obese (60.3% vs. 44.2% in matched controls, p □0.001), and this was observed both in glucocorticoid-treated and untreated CAH patients (59.1 and 61.1%, respectively). The Townsend Deprivation Index and the Charlson Comorbidity Index did not differ between CAH patients and controls.

### Glucocorticoid prescriptions

The most commonly prescribed type of glucocorticoid in the AD cohort was hydrocortisone (1296 patients, 82%), followed by prednisolone (187 patients, 11.8%). Only a minority of patients were prescribed cortisone acetate (91 patients, 5.8%, no longer available in the UK) and dexamethasone (6 patients, 0.4%).

In the glucocorticoid-treated CAH cohort, prednisolone was most commonly prescribed (127 patients, 50.0%), followed by hydrocortisone (96 patients, 37.8%), with a small minority receiving dexamethasone (15 patients, 5.9%) or cortisone acetate (11 patients, 4.3%). Only five CAH patients (2%) were prescribed a combination of short- and long-acting glucocorticoids.

### Risk of Infections

The risk of LRTIs, UTIs and GIIs was significantly increased in the AD cohort compared to unexposed patients, with the highest relative risk observed for GIIs (adjusted incidence rate ratio (aIRR) 3.80 [95% CI 2.99-4.84]) followed by LRTIs (aIRR 2.11 [95% CI 1.64-2.69]) and UTIs (aIRR 1.51 [95% CI 1.29-1.77]) (Table 3 and Figure 1). These results were confirmed in the sub-analysis of patients who had at least two prescriptions of both glucocorticoids and mineralocorticoids (94.5% of the total cohort) (Suppl. Table 1).

**Table 3:**
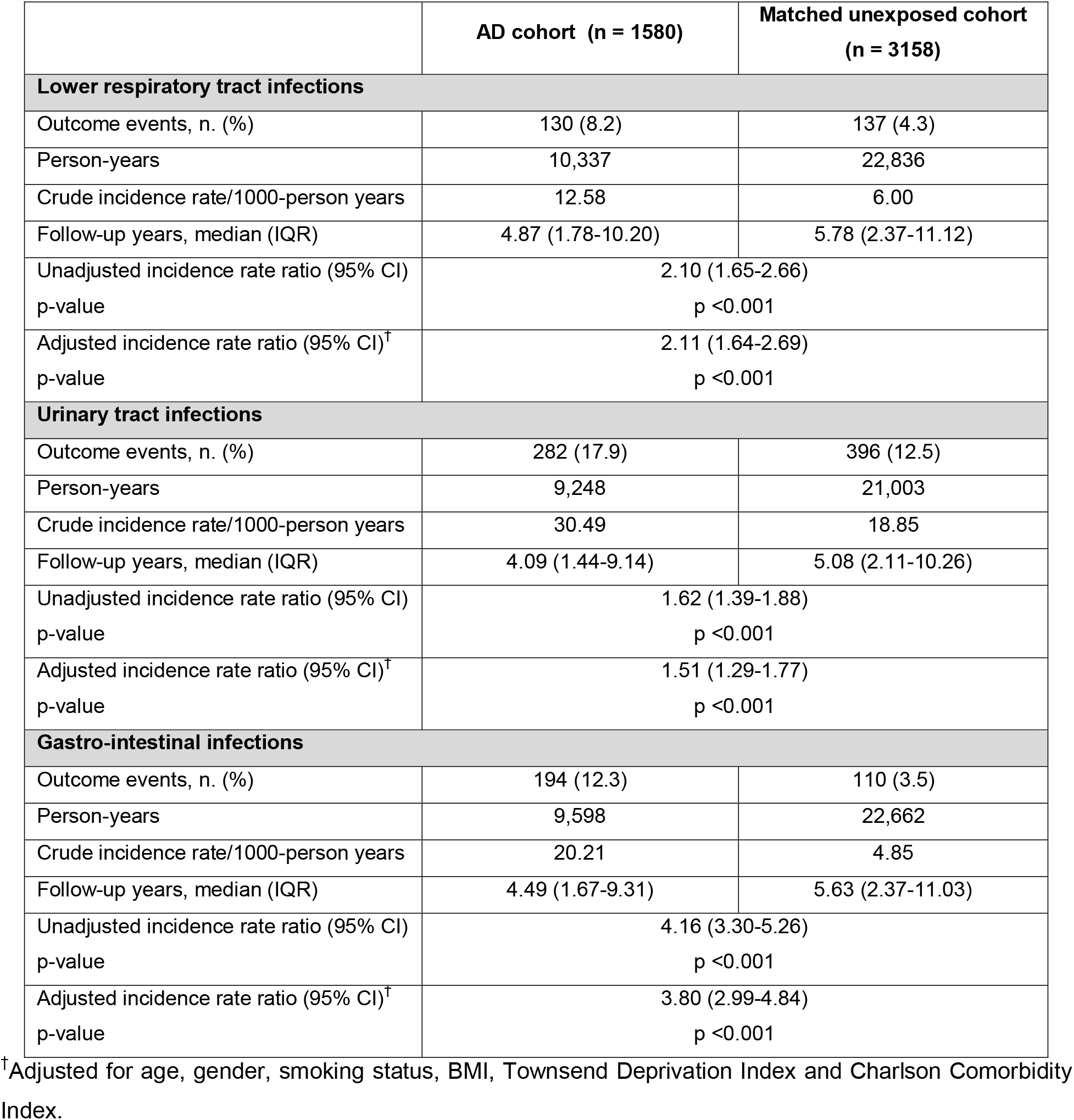
Absolute and relative risk of infections in AD patients and matched cohort.

|  | AD cohort (n = 1580) | Matched unexposed cohort (n = 3158) |
| --- | --- | --- |
| Lower respiratory tract infections |  |  |
| Outcome events, n. (%) | 130 (8.2) | 137 (4.3) |
| Person-years | 10,337 | 22,836 |
| Crude incidence rate/1000-person years | 12.58 | 6.00 |
| Follow-up years, median (IQR) | 4.87 (1.78-10.20) | 5.78 (2.37-11.12) |
| Unadjusted incidence rate ratio (95% CI) | 2.10 (1.65-2.66) |  |
| p-value | p <0.001 |  |
| Adjusted incidence rate ratio (95% CI)<sup>†</sup> | 2.11 (1.64-2.69) |  |
| p-value | p <0.001 |  |
| Urinary tract infections |  |  |
| Outcome events, n. (%) | 282 (17.9) | 396 (12.5) |
| Person-years | 9,248 | 21,003 |
| Crude incidence rate/1000-person years | 30.49 | 18.85 |
| Follow-up years, median (IQR) | 4.09 (1.44-9.14) | 5.08 (2.11-10.26) |
| Unadjusted incidence rate ratio (95% CI) | 1.62 (1.39-1.88) |  |
| p-value | p <0.001 |  |
| Adjusted incidence rate ratio (95% CI)<sup>†</sup> | 1.51 (1.29-1.77) |  |
| p-value | p <0.001 |  |
| Gastro-intestinal infections |  |  |
| Outcome events, n. (%) | 194 (12.3) | 110 (3.5) |
| Person-years | 9,598 | 22,662 |
| Crude incidence rate/1000-person years | 20.21 | 4.85 |
| Follow-up years, median (IQR) | 4.49 (1.67-9.31) | 5.63 (2.37-11.03) |
| Unadjusted incidence rate ratio (95% CI) | 4.16 (3.30-5.26) |  |
| p-value | p <0.001 |  |
| Adjusted incidence rate ratio (95% CI)<sup>†</sup> | 3.80 (2.99-4.84) |  |
| p-value | p <0.001 |  |

<sup>†</sup>Adjusted for age, gender, smoking status, BMI, Townsend Deprivation Index and Charlson Comorbidity Index.

**Figure 1 -.**
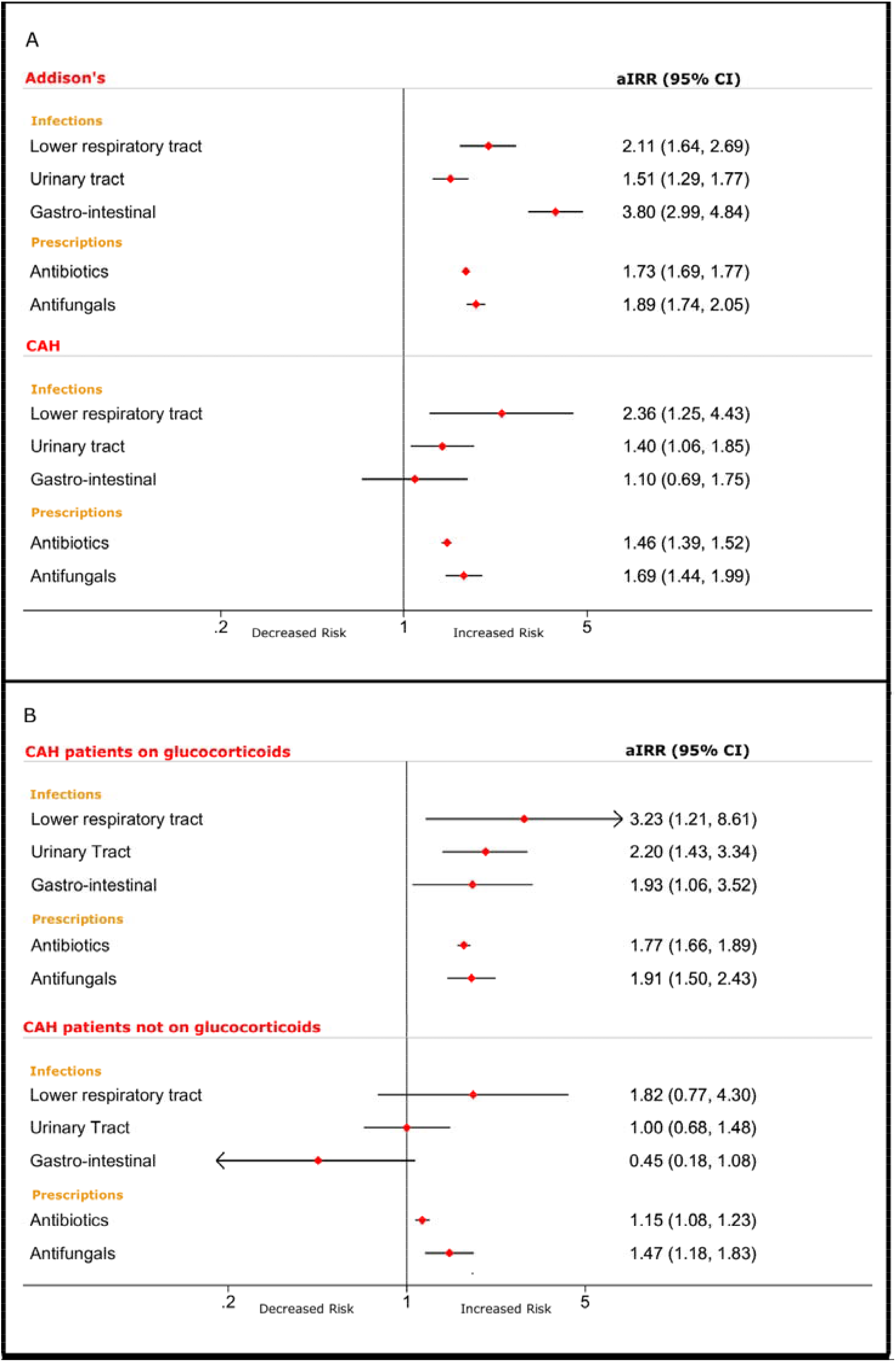
Forest plot of outcomes. Panel A: adjusted Incidence Rate Ratio (aIRR) for infections and antimicrobial prescriptions in AD and CAH cohorts. Panel B: aIRR for infections and antimicrobial prescriptions in CAH patients separately for patients with and without chronic glucocorticoid treatment.

In the overall CAH cohort, there was a significantly increased risk of UTIs and LRTIs (aIRR 1.40 [95% CI 1.06-1.85] and 2.36 [95% CI 1.25-4.42], respectively), with no difference in GI infections (Table 4 and Figure 1). However, when analyzing the population accordingly to glucocorticoid use, only patients exposed to glucocorticoids had an increased risk of infections, with the highest risk observed for LRTIs (aIRR 3.23 [95% CI 1.21-8.61]) followed by UTIs (aIRR 2.20 [95% CI 1.43-3.4]) and GIIs (aIRR 1.93 [95% CI 1.06-3.52]) (Table 4 and Figure 1). In contrast, infection risk in CAH patients not treated with glucocorticoids did not differ from that observed in the matched background population.

**Table 4:** Absolute and relative risk of infections in CAH patients and matched control cohort.

|  | CAH cohort<br>(n = 602) | Matched<br>unexposed<br>cohort (n = 1204) | CAH cohort on<br>glucocorticoids<br>(n = 254) | Matched<br>unexposed<br>cohort (n = 508) | CAH cohort not on<br>glucocorticoids<br>(n = 348) | Matched<br>unexposed<br>cohort (n = 696) |
| --- | --- | --- | --- | --- | --- | --- |
| Lower respiratory tract infections |  |  |  |  |  |  |
| Outcome events, n. (%) | 22 (3.7) | 19 (1.6) | 12 (4.7) | 7 (1.4) | 10 (2.9) | 12 (1.7) |
| Person-years | 3843 | 7842 | 1924 | 3567 | 1919 | 4275 |
| Crude incidence rate/1000-person years | 5.72 | 2.42 | 6.24 | 1.96 | 5.21 | 2.81 |
| Follow-up years, median (IQR) | 4.83 (1.92-9.54) | 5.10 (2.04-9.80) | 6.00 (2.60-12.06) | 5.17 (2.45-10.77) | 4.07 (1.54-8.00) | 4.93 (1.86-9.37) |
| Unadjusted incidence rate ratio (95% CI) | 2.36 (1.28-4.36) |  | 3.18 (1.25-8.07) |  | 1.86 (0.80-4.30) |  |
| p-value | p = 0.01 |  | p = 0.02 |  | p = 0.15 |  |
| Adjusted incidence rate ratio (95% CI)<sup>†</sup> | 2.36 (1.25-4.43) |  | 3.23 (1.21-8.61) |  | 1.82 (0.77-4.30) |  |
| p-value | p = 0.01 |  | p = 0.02 |  | p = 0.17 |  |
| Urinary tract infections |  |  |  |  |  |  |
| Outcome events, n. (%) | 83 (13.8) | 130 (10.8) | 45 (17.7) | 43 (8.5) | 38 (10.9) | 87 (12.5) |
| Person-years | 3478 | 7217 | 1709 | 3374 | 1769 | 3843 |
| Crude incidence rate/1000-person years | 23.87 | 18.01 | 26.33 | 12.75 | 21.48 | 22.64 |
| Follow-up years, median (IQR) | 4.15 (1.59-8.80) | 4.64 (1.83-8.80) | 4.97 (2.09-9.77) | 4.84 (2.22-9.77) | 3.42 (1.24-7.53) | 4.18 (1.69-7.90) |
| Unadjusted incidence rate ratio (95% CI) | 1.32 (1.01-1.74) |  | 2.07 (1.36-3.14) |  | 0.95 (0.65-1.39) |  |
| p-value | p = 0.05 |  | p <0.001 |  | p = 0.79 |  |
| Adjusted incidence rate ratio (95% CI)<sup>†</sup> | 1.40 (1.06-1.85) |  | 2.20 (1.43-3.34) |  | 1.00 (0.68-1.48) |  |
| p-value | p = 0.02 |  | p <0.001 |  | p = 0.99 |  |
| Gastro-intestinal infections |  |  |  |  |  |  |
| Outcome events, n. (%) | 29 (4.8) | 52 (4.3) | 23 (9.1) | 23 (4.5) | 6 (1.7) | 29 (4.2) |
| Person-years. | 3773 | 7678 | 1825 | 3486 | 1948 | 4192 |
| Crude incidence rate/1000-person years | 7.69 | 6.77 | 12.60 | 6.60 | 3.08 | 6.92 |
| Follow-up years, median (IQR) | 4.70 (1.83-9.63) | 4.94 (2.04-9.69) | 5.82 (2.12-11.38) | 5.14 (2.47-10.38) | 4.09 (1.50-8.11) | 4.79 (1.86-9.13) |
| Unadjusted incidence rate ratio (95% CI) | 1.13 (0.72-1.79) |  | 1.91 (1.07-3.40) |  | 0.45 (0.18-1.07) |  |
| p-value | p = 0.59 |  | p = 0.03 |  | p = 0.07 |  |
| Adjusted incidence rate ratio (95% CI)<sup>†</sup> | 1.10 (0.69-1.75) |  | 1.93 (1.06-3.52) |  | 0.45 (0.18-1.08) |  |
| p-value | p = 0.70 |  | p = 0.03 |  | p = 0.07 |  |

485 <sup>†</sup>Adjusted for age, gender, smoking status, BMI, Townsend Deprivation Index and Charlson Comorbidity Index.

### Antimicrobial prescriptions

Prescription rates of antibiotics and antifungals were increased in patients with AD (aIRR 1.73 [95% CI 1.69-1.77] and 1.89 [95% CI 1.74-2.05], respectively) (Table 5 and Figure 1). These results were confirmed in the sub-analysis of patients who had at least two prescriptions of both glucocorticoids and mineralocorticoids (Suppl. Table 2).

**Table 5:** Antimicrobial prescriptions counts in AD patients compared to the matched control cohort.

|  | AD cohort (n = 1580) | Matched unexposed cohort (n = 3158) |
| --- | --- | --- |
| Antibiotic prescriptions |  |  |
| Count of prescriptions, n. | 13286 | 15884 |
| Person-years | 10767 | 23308 |
| Count rates (per 1000 years) | 1234 | 681 |
| Follow-up years, median (IQR) | 5.12 (1.95-10.77) | 5.90 (2.54-11.33) |
| Unadjusted incidence rate ratio (95% CI) | 1.81 (1.77-1.85) |  |
| p-value | p <0.001 |  |
| Adjusted incidence rate ratio (95% CI)<sup>†</sup> | 1.73 (1.69-1.77) |  |
| p-value | p <0.001 |  |
| Antifungal prescriptions |  |  |
| Count of prescriptions, n. | 1191 | 1213 |
| Person-years | 10767 | 23308 |
| Count rates (per 1000 years) | 111 | 52 |
| Follow-up years, median (IQR) | 5.12 (1.95-10.77) | 5.90 (2.54-11.33) |
| Unadjusted incidence rate ratio (95% CI) | 2.13 (1.96-2.30) |  |
| p-value | p <0.001 |  |
| Adjusted incidence rate ratio (95% CI)<sup>†</sup> | 1.89 (1.74-2.05) |  |
| p-value | p <0.001 |  |

<sup>†</sup>Adjusted for age, gender, smoking status, BMI, Townsend Deprivation Index and Charlson Comorbidity Index.

Similarly, we observed increased antimicrobial prescription rates in CAH patients, with a higher prescription rate in glucocorticoid-treated patients (antibiotics: aIRR 1.77 [95% CI 1.66-1.89]; antifungals: aIRR 1.91 [95% CI 1.50-2.43]) than in CAH patients not exposed to glucocorticoids (antibiotics: aIRR 1.15 [95% CI 1.08-1.23]; antifungals: aIRR 1.44 [95% CI 1.18-1.83]) (Table 6 and Figure 1).

**Table 6:** Antimicrobial prescription counts in CAH patients compared to the matched control cohort.

|  | CAH cohort<br>(n = 602) | Matched<br>unexposed<br>cohort (n = 1204) | CAH cohort on<br>glucocorticoids<br>(n = 254) | Matched<br>unexposed<br>cohort (n = 508) | CAH cohort not on<br>glucocorticoids<br>(n = 348) | Matched<br>unexposed<br>cohort (n = 696) |
| --- | --- | --- | --- | --- | --- | --- |
| Antibiotics prescriptions |  |  |  |  |  |  |
| Count of prescriptions, n. | 3543 | 4930 | 2134 | 2088 | 1409 | 2842 |
| Person-years | 3941 | 7957 | 1974 | 3611 | 1967 | 4346 |
| Count rates (per 1000 years) | 899 | 619 | 1081 | 578 | 716 | 654 |
| Follow-up years, median (IQR) | 4.85 (1.95-10.32) | 5.23 (2.09-9.90) | 6.12 (2.67-12.48) | 5.32 (2.54-10.87) | 4.12 (1.54-8.30) | 5.20 (1.88-9.62) |
| Unadjusted incidence rate ratio (95% CI) | 1.45 (1.39-1.51) |  | 1.87 (1.76-1.99) |  | 1.10 (1.03-1.17) |  |
| p-value | p <0.001 |  | p <0.001 |  | p = 0.01 |  |
| Adjusted incidence rate ratio (95% CI)<sup>†</sup> | 1.46 (1.39-1.52) |  | 1.77 (1.66-1.89) |  | 1.15 (1.08-1.23) |  |
| p-value | p <0.001 |  | p <0.001 |  | p <0.001 |  |
| Antifungals prescriptions |  |  |  |  |  |  |
| Count of prescriptions, n. | 302 | 330 | 159 | 130 | 143 | 200 |
| Person-years | 3941 | 7957 | 1974 | 3611 | 1967 | 4346 |
| Count rates (per 1000 years) | 77 | 41 | 81 | 36 | 73 | 46 |
| Follow-up years, median (IQR) | 4.85 (1.95-10.32) | 5.23 (2.09-9.90) | 6.12 (2.67-12.48) | 5.32 (2.54-10.87) | 4.12 (1.54-8.30) | 5.20 (1.88-9.62) |
| Unadjusted incidence rate ratio (95% CI) | 1.85 (1.58-2.16) |  | 2.24 (1.77-2.82) |  | 1.58 (1.27-1.96) |  |
| p-value | p <0.001 |  | p <0.001 |  | p <0.001 |  |
| Adjusted incidence rate ratio (95% CI)<sup>†</sup> | 1.69 (1.44-1.99) |  | 1.91 (1.50-2.43) |  | 1.47 (1.18-1.83) |  |
| p-value | p <0.001 |  | p <0.001 |  | p <0.001 |  |

<sup>†</sup>Adjusted for age, gender, smoking status, BMI, Townsend Deprivation Index and Charlson Comorbidity Index.

Given the higher incidence of type 1 diabetes mellitus (T1DM) in our AD cohort (8% vs. 0.5% in matched controls), and given the potentially higher risk of infections in T1DM patients, we performed a sub group analysis comparing AD patients with and without T1DM to matched unexposed cohort. Findings were similar, though given the smaller number of T1DM patients group, some did not reach statistical significance (Suppl. Tables 3 and 4).

## DISCUSSION

In this population-based study we found that the risk of three common infections (lower respiratory tract, urinary tract, and gastrointestinal infections) was increased in the primary care setting in patients with PAI, as compared to population-based matched controls. This was also supported by our finding of increased prescription rates of antimicrobials in in patients with PAI. Moreover, we found that CAH patients not receiving glucocorticoids did not have an increased risk of infections, indicating that glucocorticoid therapy might at least partly drive the increased infection risk observed in PAI. To our knowledge, our study is the first to analyze the risk of infection in PAI according to different etiologies and also the first to evaluate these outcomes in a primary care setting.

Previous studies have described an increased infection-related mortality in patients with AD (2,4), but not in CAH patients (3). This was attributed to infections representing a possible trigger for a fatal adrenal crisis. Smans and colleagues reported an increase of the use of antimicrobials and of infection-related hospital admissions in PAI (5); however, the authors focused on hospital-treated infections only, possibly overestimating the actual incidence of this complication due to a lower threshold for admission in PAI patients. In addition, information on the actual etiology of PAI was not available in this study, as PAI was diagnosed based on concomitant glucocorticoid and mineralocorticoid prescriptions, which did not allow to differentiate between AD, CAH, and other causes of PAI.

Until recently, it was unclear whether the observed increase in infection episodes in patients with PAI is related to the underlying disease itself or to the non-physiological delivery of glucocorticoid replacement by currently available glucocorticoid preparations. Autoimmune AD patients frequently also suffer from other autoimmune comorbidities (18), and this was confirmed in our study, with more prevalent autoimmune disease in our AD cohort, which can be safely assumed to consist of a large majority of patients with AD of autoimmune origin. However, in CAH patients, there is only marginal evidence of an imbalance of immune function (19), and as we found similar increases in infection risk in the CAH cohort, potential etiology-related immune function is unlikely to explain the increased susceptibility to infections we observed.

Supraphysiological glucocorticoid doses, as usually administered in the context of chronic inflammatory disease, is well known to cause changes in the immune system, with consequently increased risk of bacterial and fungal infections (20). However, this has not been demonstrated for the physiological replacement doses generally used in patients with PAI. Still, currently available glucocorticoid replacement therapy does not provide a physiological substitution, with significant peaks and troughs of cortisol availability during the day following oral intake of immediate release glucocorticoid preparations. In addition, significant heterogeneity exists in the management of glucocorticoid replacement in clinical practice; a recent paper recorded 25 different regimens with which glucocorticoid therapy is administered in AD patients receiving a daily hydrocortisone of 20 mg (21). Therefore, it would come as no surprise that also physiological dose glucocorticoid therapy is not free of side effects, if administered in a non-physiological delivery pattern. An improvement in metabolic outcomes after switching from standard cortisol replacement to more physiological cortisol replacement via continuous subcutaneous hydrocortisone was previously demonstrated in both AD and CAH patients (22,23).

Some recent papers have indeed suggested that adverse changes in immune function might occur with glucocorticoid replacement in PAI. A recent paper documented significantly decreased natural killer cytotoxicity in patients with PAI (7), which was present in both patients with autoimmune adrenalitis and those with PAI following bilateral adrenalectomy, indicating that the underlying etiology did not play a role in these changes in immune function. A recent randomized control trial including patients with primary and secondary adrenal insufficiency reported a reduction in respiratory tract infections with modified-release hydrocortisone (8).

However, this was a secondary outcome, based on self-reported questionnaires on infections and not verified against medical records, thereby providing only limited evidence. A study on immune function in the same cohort reported dysregulation of circadian gene expression in peripheral blood mononuclear cells in the PAI patients at baseline, which attenuated after the switch to modified-release hydrocortisone therapy (24). The findings of our study, including both patients with AD and CAH, suggest that exogenous glucocorticoid is at least a contributory factor to the increased infection risk we observed, given that no significant increase in infection risk was observed in the CAH patients not receiving glucocorticoid therapy.

Both our AD and CAH populations had increased prescription rates for antibiotics and antifungals. Interestingly, increased prescription rates were also noted in the CAH patients not receiving glucocorticoid treatment, albeit to a much lower extent. This could possibly be explained by a lower threshold for prescribing antimicrobials due to the perceived risk of adrenal crisis in CAH patients; in fact, up to 60% of non-classic CAH patients, who usually do not receive chronic glucocorticoid replacement, have been reported to have at least partial glucocorticoid deficiency as assessed by cosyntropin testing (25).

The highest increase in risk of infection in our AD cohort was seen in GIIs, while for the CAH cohort on glucocorticoids the most significant increase in risk was seen in LRTIs and UTIs; however, the differences between the three infection groups was not statistically significant. This may be explained by the age difference between AD and CAH patients, with mean ages of 51.7 and 35.4 years, respectively. Indeed, LRTIs and UTIs are more frequently diagnosed in older patients (26,27), and this was noted in our matched populations as well (population matched for AD: LRTIs 4.3%, UTIs 12.5%; population matched for CAH patients on glucocorticoids: LRTIs 1.4%, UTIs 8.5%). Therefore, the higher aIRR of LRTIs and UTIs in CAH patients is probably related to a difference in age-related background risk.

Our AD cohort had an age and sex distribution similar to the one reported in other papers (2,5), and the types of prescribed glucocorticoid preparations at baseline in this cohort were not different from the ones reported in a recent worldwide survey (28). Our CAH cohort was younger that the AD cohort, consistent with the different etiology of these two diseases, and the types of glucocorticoids prescribed was similar to those reported in the cross-sectional UK CaHASE study (29), with the possible exception of lower numbers of dexamethasone users in our study. Taking this into account, our results can be assumed to be representative of the UK AD and CAH populations.

Our study has several strengths. We used a large population-based sample of patients of both sexes, across all adult age groups, with very strict inclusion and exclusion criteria, allowing us to include only patients with a true diagnosis of AD and CAH. Using the cohort study design, allowed us to look at longitudinal occurrence of infections and antimicrobial use. There are also some limitations to our study. Firstly, all data relies on the accurate recording of diagnoses by GPs and this could have resulted in some degree of misclassification of the exposed cohorts and of the different episodes of infection. Though general practitioners document reasons for consultations in the electronic medical records, it is possible that when a patient presented with two or more conditions this may have not been accurately coded; however all prescriptions are electronically documented and therefore are captured accurately. Secondly, the threshold for visiting GP might be lower in patients with PAI who receive regular education on the importance of treating infections promptly to avoid adrenal crisis. This may be a factor resulting in a degree of overestimation of the difference in the infection rates we found between these cohorts. However, since patients with PAI are generally more medicalized, it is also possible that they own a higher knowledge of diseases and might decide to treat themselves without consulting the GP. Thirdly, we could not evaluate the influence of different doses or types of glucocorticoid on the outcomes of interest due to the methodology used and due to the small number of infection events when further subdividing our populations according to type of glucocorticoid. Furthermore, even though we tried to assess the impact of associated comorbidities by adjusting for Charlson Comorbidity Index, this does not exclude the possibility that some other confounders not accounted for in our analyses might have influenced our results. Lastly, though there is some evidence of an immune-modulatory effect of androgens (30), we could not take this into account in our population as we had no data on DHEA replacement therapy in AD patients, since this is a hospital-prescribed drug in the UK; similarly, we did not have data on biochemical control of androgen excess in the CAH patients.

Our findings have several practical implications. Firstly, given the confirmation of a higher risk of infections in patients with PAI due to AD and CAH, all healthcare professionals involved in the care of PAI patients should have a heightened alertness for the possibility of infections in these patients. This may also provide a case for recommending a vaccination strategy in PAI, e.g. against *Streptococcus pneumoniae*, the leading cause of LRTIs in adults (31), in order to reduce the risk of these infections and related morbidity and mortality. Secondly, our paper provides additional evidence that non-physiological delivery of glucocorticoid replacement by currently available preparations represents a risk factor for the development of infections. This supports the case for a therapeutic shift towards more physiological replacement therapy options in these patients (32). Future studies will have to clarify whether achieving a more physiological delivery of glucocorticoid replacement will decrease the risk of infections in PAI, with the potential to result in reduced morbidity and mortality in these patients.

## Supporting information

Suppl Material incl. Suppl. Tables 1-4

## ACKNOWLEDGMENTS

This work was supported by the Medical Research Council UK (Program Grant 0900567, to W.A.). K.N. is a UK Research and Innovation (UKRI)/Health Data Research (HDR) UK Innovation Clinical Fellow. W.A. receives support from the National Institute of Health Research (NIHR) Birmingham Biomedical Research Centre (BRC-1215-20009). The views expressed in this publication are those of the authors and not necessarily those of the NIHR or the Department of Health and Social Care UK.

