## Supplementary material for "Increased infection risk in Addison’s disease and congenital adrenal hyperplasia: a primary care database cohort study": Suppl Material incl. Suppl. Tables 1-4

**Supplemental Material	Tresoldi et al.	doi:** https://doi.org/10.1101/628156

**Appendix – Read codes for specific outcomes**

Codes for diagnosis of Addison’s Disease (AD; non-CAH primary adrenal insufficiency)

Medcode, description, freq, rank

C154100,Addison's disease,4698,1

C154600,Addisonian crisis,543,1

C154000,Acute adrenal insufficiency,194,1

C154.00,Corticoadrenal insufficiency,526,1

C154011,Addisonian crisis,174,1

C154012,Adrenal crisis,48,1

C154z00,Corticoadrenal insufficiency NOS,83,1

C154z11,Adrenal hypofunction,66,1

C154z12,Adrenal insufficiency NEC,359,1

C155.00,Other adrenal hypofunction,25,1

C155z00,Other adrenal hypofunction NOS,23,1

Codes for diagnosis of congenital adrenal hyperplasia (CAH)

CODE,description

C152000,Congenital adrenogenital syndrome

C152200,Defective synthesis of 21 hydroxylase

C152300,Defective synthesis of 11B hydroxylase

C152400,Defective synthesis of 3B hydroxysteroid dehydrogenase

C152500,Defective synthesis of 17-20 desmolase

C152600,Defective synthesis of 17 alpha hydroxylase

C152700,Other adrenogenital syndrome with salt loss

C152800,Other adrenogenital syndrome without mention of salt loss

C152811,Adrenogenital syndrome NOS

C152812,Congenital adrenal hyperplasia NEC

C152900,Precocious puberty with adrenocortical hyperfunction

C152A11,Virilisation - adrenogenital

C152.00,Adrenogenital disorders

C152912,Precocious puberty with adrenal hyperplasia

Codes for Glucocorticoid (GC) prescriptions

drugcode,genericname,FLAG

84044998,Hydrocortisone 10mg/5ml oral suspension,2

86221979,Hydrocortisone sodium succinate 100mg powder for solution for injection vials,2

86220979,Hydrocortisone sodium succinate 100mg powder and solvent for solution for injection vials,2

85251998,Hydrocortisone sodium phosphate 100mg/1ml solution for injection ampoules,2

85249998,Hydrocortisone sodium phosphate 100mg/1ml solution for injection ampoules,2

92412998,Hydrocortisone 5mg/5ml oral solution,2

96680992,Hydrocortisone sodium succinate 100mg powder for solution for injection vials,2

96143997,Hydrocortisone 20mg tablets,2

96143998,Hydrocortisone 10mg tablets,2

96172998,Hydrocortisone sodium phosphate 100mg/1ml solution for injection ampoules,2

96175998,Hydrocortisone sodium succinate 100mg powder for solution for injection vials,2

96270992,Hydrocortisone sodium succinate 100mg powder for solution for injection vials,2

97341992,Efcortelan 100 mg inj,2

95833992,Hydrocortisone sodium succinate pow,2

97583992,Hydrocortisone 25 mg tab,2

97640998,Hydrocortisone sodium succinate 100mg powder for solution for injection vials,2

99550998,Hydrocortisone 20mg tablets,2

98376998,Hydrocortisone na succinate 100mg/vial injection,2

99896992,Hydrocortisone sodium phosphate 5 mg sol,2

97492998,Hydrocortisone 10mg tablets,2

97492997,Hydrocortisone 20mg tablets,2

53807979,Hydrocortisone 5mg modified-release tablets,2

53808979,Hydrocortisone 5mg modified-release tablets,2

53809979,Hydrocortisone 20mg modified-release tablets,2

64761979,Hydrocortisone sodium succinate 100mg powder and solvent for solution for injection vials,2

53810979,Hydrocortisone 20mg modified-release tablets,2

64762979,Hydrocortisone sodium succinate 100mg powder for solution for injection vials,2

94471998,Hydrocortisone acetate 25mg/1ml suspension for injection ampoules,2

98252990,Hydrocortisone acetate powder,2

99549998,Hydrocortisone acetate 25mg/1ml suspension for injection ampoules,2

87950998,Hydrocortisone 2.5mg muco-adhesive buccal tablets sugar free,2

91055979,Hydrocortisone 2.5mg muco-adhesive buccal tablets sugar free,2

92046990,Hydrocortisone 2.5mg muco-adhesive buccal tablets sugar free,2

92064990,Hydrocortisone 2.5mg muco-adhesive buccal tablets sugar free,2

93588998,Hydrocortisone 2.5mg muco-adhesive buccal tablets sugar free,2

96269992,Hydrocortisone 2.5mg muco-adhesive buccal tablets sugar free,2

99806998,Hydrocortisone 2.5mg lozenges,2

79760979,Hydrocortisone 5mg/5ml oral suspension,2

63230979,Hydrocortisone 100mg suppositories,2

59794979,Hydrocortisone 5mg/5ml oral suspension sugar free,2

67795979,Hydrocortisone 3mg/5ml oral suspension,2

63726979,Hydrocortisone 2mg capsules,2

63611979,Hydrocortisone 2.5mg capsules,2

99184989,Hydrocortisone 25mg suppositories,2

97064998,Hydrocortisone 25mg suppositories,2

85117998,Hydrocortisone tablet,2

98804990,Hydrocortisone powder,2

83566978,Prednisolone 2.5mg gastro-resistant tablets,3

91648990,Prednisolone 2.5mg gastro-resistant tablets,3

93075998,Prednisolone 5mg soluble tablets,3

93369979,Prednisolone 5mg soluble tablets,3

93887990,Prednisolone 5mg gastro-resistant tablets,3

93370979,Prednisolone 5mg soluble tablets,3

93912997,Prednisolone 5mg gastro-resistant tablets,3

93912998,Prednisolone 2.5mg gastro-resistant tablets,3

91788990,Prednisolone 5mg soluble tablets,3

97155997,Prednisolone 5mg gastro-resistant tablets,3

96577990,Prednisolone 5mg soluble tablets,3

95417998,Prednisolone 2.5mg tablets,3

95417996,Prednisolone 5mg gastro-resistant tablets,3

95912990,Prednisolone 25mg tablets,3

97101990,Prednisolone 2.5mg gastro-resistant tablets,3

96411992,Prednisolone 15 mg tab,3

97101989,Prednisolone 5mg gastro-resistant tablets,3

95594990,Prednisolone 1mg tablets,3

95593990,Prednisolone 5mg gastro-resistant tablets,3

97240992,Prednisolone 1mg tablets,3

97155998,Prednisolone 1mg tablets,3

96744992,Prednisolone 4 mg tab,3

95493992,Prednisolone 2 mg tab,3

95492992,Prednisolone 10 mg tab,3

95487992,Prednisolone 5mg soluble tablets,3

95484992,Prednisolone 1mg tablets,3

96361990,Prednisolone 1mg tablets,3

96361989,Prednisolone 5mg gastro-resistant tablets,3

99424989,Prednisolone 5mg gastro-resistant tablets,3

99423989,Prednisolone 5mg gastro-resistant tablets,3

99423988,Prednisolone 2.5mg gastro-resistant tablets,3

97726990,Prednisolone 2.5mg gastro-resistant tablets,3

99226998,Prednisolone 5mg soluble tablets,3

99425990,Prednisolone 1mg tablets,3

98562997,Prednisolone 5mg gastro-resistant tablets,3

98107990,Prednisolone 5mg gastro-resistant tablets,3

98562998,Prednisolone 2.5mg gastro-resistant tablets,3

98455990,Prednisolone 5mg gastro-resistant tablets,3

99228998,Prednisolone 1mg tablets,3

99228997,Prednisolone 5mg tablets,3

98514997,Prednisolone 5mg tablets,3

98514998,Prednisolone 1mg tablets,3

99100990,Prednisolone 1mg tablets,3

99100989,Prednisolone 5mg gastro-resistant tablets,3

99099990,Prednisolone 2.5mg gastro-resistant tablets,3

99099989,Prednisolone 5mg gastro-resistant tablets,3

99099988,Prednisolone 5mg gastro-resistant tablets,3

99425988,Prednisolone 5mg gastro-resistant tablets,3

99424990,Prednisolone 1mg tablets,3

99137998,Prednisolone 5mg tablets,3

97929990,Prednisolone 5mg gastro-resistant tablets,3

99423990,Prednisolone 1mg tablets,3

99425989,Prednisolone 5mg gastro-resistant tablets,3

98456989,Prednisolone 5mg gastro-resistant tablets,3

98456988,Prednisolone 1mg tablets,3

53396978,Prednisolone 2.5mg tablets,3

53398978,Prednisolone 10mg tablets,3

97237992,Deltastab 2 mg tab,3

88912979,Prednisolone 10mg/5ml oral solution,3

66092979,Prednisolone 2.5mg/5ml oral suspension,3

66096979,Prednisolone 1mg/5ml oral suspension,3

65924979,Prednisolone 5mg/5ml oral solution,3

81915998,Prednisone 2mg modified-release tablets,4

81916998,Prednisone 1mg modified-release tablets,4

81911998,Prednisone 5mg modified-release tablets,4

81914998,Prednisone 5mg modified-release tablets,4

83565978,Prednisolone 5mg gastro-resistant tablets,4

81913998,Prednisone 1mg modified-release tablets,4

81912998,Prednisone 2mg modified-release tablets,4

96743992,Prednisone 2.5 mg tab,4

97156997,Prednisone 5mg tablets,4

97156998,Prednisone 1mg tablets,4

97942992,Prednisone 10 mg tab,4

99098990,Prednisone 5mg tablets,4

99781998,Prednisone 5mg tablets,4

87704998,Cortisone 5mg capsules,5

94442992,Cortisone 25mg tablets,5

96603998,Cortisone acetate 5mg tablets,5

96603997,Cortisone 25mg tablets,5

97203992,Cortisone acetate 2.5 mg tab,5

94870992,Cortisone acetate msd 5 mg tab,5

97204992,Cortisone acetate 25 mg inj,5

99802998,Cortisone acetate 25mg tablets,5

99803997,Cortisone acetate 25mg tablets,5

99803998,Cortisone acetate 5mg tablets,5

99804998,Cortisone acetate 25mg tablets,5

83825998,Dexamethasone 500microgram tablets,6

93423979,Dexamethasone 2mg/5ml oral solution,6

93427979,Dexamethasone 2mg tablets,6

93429979,Dexamethasone 2mg tablets,6

93431979,Dexamethasone 500microgram tablets,6

91646998,Dexamethasone 2mg/5ml oral solution sugar free,6

92810997,Dexamethasone 2mg/5ml oral solution sugar free,6

92810998,Dexamethasone 2mg/5ml oral solution,6

93098990,Dexamethasone 2mg tablets,6

92484990,Dexamethasone 500microgram tablets,6

93867997,Dexamethasone 100mg/5ml solution for injection vials,6

94908992,Dexacortisyl 2 mg tab,6

95357992,Dexamethasone 500mcg tablets,6

96182992,Dexamethasone 750 mcg tab,6

97243992,Dexamethasone 2mg tablets,6

96431997,Dexamethasone 2mg tablets,6

96431996,Dexamethasone 500micrograms/5ml oral solution,6

98724997,Dexamethasone 2mg tablets,6

98644990,Dexamethasone 500microgram tablets,6

98072989,Dexamethasone 500micrograms/5ml oral solution,6

98072990,Dexamethasone 2mg/5ml oral solution,6

98644989,Dexamethasone 2mg tablets,6

97502998,Dexamethasone 500mcg tablets,6

97493998,Dexamethasone 100mg/5ml injection,6

70200979,Dexamethasone 2mg/5ml oral solution sugar free,6

70202979,Dexamethasone 2mg/5ml oral solution,6

52504979,Dexamethasone 2mg/5ml oral solution,6

80402979,Dexamethasone 1mg/5ml oral solution,6

70228979,Dexamethasone 1.5mg/5ml oral solution,6

85515998,Dexamethasone 100microgram capsules,6

Codes for mineralocorticoid (MC) prescriptions

drugcode,genericname,FLAG

96526998,Fludrocortisone 100microgram tablets,1

96658992,Fludrocortisone 25 mcg tab,1

97445992,Florinef .2 mg tab,1

95026992,Fludrocortisone 100microgram tablets,1

96245992,Fludrocortisone .05 mg sus,1

97452992,Fludrocortisone 20 mcg tab,1

97451992,Fludrocortisone 75 mcg tab,1

99636998,Fludrocortisone 100microgram tablets,1

80070979,Fludrocortisone 50micrograms/5ml oral suspension,1

80076979,Fludrocortisone 25micrograms/5ml oral suspension,1

69470979,Fludrocortisone 20micrograms/5ml oral suspension,1

69468979,Fludrocortisone 30micrograms/5ml oral suspension,1

79112979,Fludrocortisone 50microgram capsules,1

84906998,Fludrocortisone liquid,1

84905998,Fludrocortisone capsule,1

Codes for Lower Respiratory Tract Infections (LRTIs)

medcode, description

H062.00,Acute lower respiratory tract infection

H06z100,Lower resp tract infection

H06z112,Acute lower respiratory tract infection

H21..00,Lobar (pneumococcal) pneumonia

H22yz00,Pneumonia due to bacteria NOS

H22z.00,Bacterial pneumonia NOS

H25..00,Bronchopneumonia due to unspecified organism

H25..11,Chest infection - unspecified bronchopneumonia

H260.00,Lobar pneumonia due to unspecified organism

H261.00,Basal pneumonia due to unspecified organism

H2B..00,Community acquired pneumonia

H2C..00,Hospital acquired pneumonia

Hyu1.00,[X]Other acute lower respiratory infections

A3BX400,Streptococ pneumon/cause/disease classified/oth chapters

A3BXB00,Klebsiella pneumoniae/cause/disease classifd/oth chapters

AyuKA00,[X]Klebsiella pneumoniae/cause/disease classifd/oth chapters

H21..11,Chest infection - pneumococcal pneumonia

H22..00,Other bacterial pneumonia

H22..11,Chest infection - other bacterial pneumonia

H220.00,Pneumonia due to klebsiella pneumoniae

H221.00,Pneumonia due to pseudomonas

H222.00,Pneumonia due to haemophilus influenzae

H222.11,Pneumonia due to haemophilus influenzae

H223.00,Pneumonia due to streptococcus

H223000,"Pneumonia due to streptococcus, group B"

H224.00,Pneumonia due to staphylococcus

H22y.00,Pneumonia due to other specified bacteria

H22y000,Pneumonia due to escherichia coli

H22y011,E.coli pneumonia

H22yX00,Pneumonia due to other aerobic gram-negative bacteria

H232.00,Pneumonia due to pleuropneumonia like organisms

AyuK300,[X]Streptococ pneumon/cause/disease classified/oth chapters

H060600,Acute pneumococcal bronchitis

H23..00,Pneumonia due to other specified organisms

H23..11,Chest infection - pneumonia organism OS

H26..00,Pneumonia due to unspecified organism

Hyu0900,[X]Pneumonia due to other aerobic gram-negative bacteria

Hyu0A00,[X]Other bacterial pneumonia

Hyu0C00,[X]Pneumonia in bacterial diseases classified elsewhere

Hyu0H00,"[X]Other pneumonia, organism unspecified"

SP13100,Other aspiration pneumonia as a complication of care

H2...00,Pneumonia and influenza

H20..00,Viral pneumonia

H20..11,Chest infection - viral pneumonia

H200.00,Pneumonia due to adenovirus

H201.00,Pneumonia due to respiratory syncytial virus

H202.00,Pneumonia due to parainfluenza virus

H203.00,Pneumonia due to human metapneumovirus

H20y.00,Viral pneumonia NEC

H20y000,Severe acute respiratory syndrome

H20z.00,Viral pneumonia NOS

H22y100,Pneumonia due to proteus

H22y200,Pneumonia - Legionella

H22y300,Pneumonia due to Gram neg bact

H230.00,Pneumonia due to Eaton's agent

H231.00,Pneumonia due to mycoplasma pneumoniae

H233.00,Chlamydial pneumonia

H23z.00,Pneumonia due to specified organism NOS

H24..00,Pneumonia with infectious diseases EC

H24..11,Chest infection with infectious disease EC

H240.00,Pneumonia with measles

H241.00,Pneumonia with cytomegalic inclusion disease

H242.00,Pneumonia with ornithosis

H243.00,Pneumonia with whooping cough

H243.11,Pneumonia with pertussis

H244.00,Pneumonia with tularaemia

H245.00,Pneumonia with anthrax

H246.00,Pneumonia with aspergillosis

H247.00,Pneumonia with other systemic mycoses

H247000,Pneumonia with candidiasis

H247100,Pneumonia with coccidioidomycosis

H247200,Pneumonia with histoplasmosis

H247z00,Pneumonia with systemic mycosis NOS

H24y.00,Pneumonia with other infectious diseases EC

H24y000,Pneumonia with actinomycosis

H24y100,Pneumonia with nocardiasis

H24y200,Pneumonia with pneumocystis carinii

H24y300,Pneumonia with Q-fever

H24y400,Pneumonia with salmonellosis

H24y500,Pneumonia with toxoplasmosis

H24y600,Pneumonia with typhoid fever

H24y700,Pneumonia with varicella

H24yz00,Pneumonia with other infectious diseases EC NOS

H24z.00,Pneumonia with infectious diseases EC NOS

H26..11,Chest infection - pnemonia due to unspecified organism

H260000,Lung consolidation

H262.00,Postoperative pneumonia

H263.00,"Pneumonitis, unspecified"

H270.00,Influenza with pneumonia

H270.11,Chest infection - influenza with pneumonia

H270000,Influenza with bronchopneumonia

H270100,"Influenza with pneumonia, influenza virus identified"

H270z00,Influenza with pneumonia NOS

H28..00,Atypical pneumonia

H2F0.00,Inflnza wth pmn due sesnl vir

H2y..00,Other specified pneumonia or influenza

H2z..00,Pneumonia or influenza NOS

Codes for Urinary Tract Infections (UTIs)

ReadCodes, Description

1A55.00,Dysuria

46B3.00,Urine bacteria test: positive

46f2.00,Urine leucocyte test = +

46f3.00,Urine leucocyte test = ++

46f4.00,Urine leucocyte test = +++

46f6.00,Urine leucocyte test = ++++

46G4.11,Leucocytes in urine

46G4.11,Leucocytes in urine

46U2.00,Urine culture - mixed growth

46U3.00,Urine culture - E. Coli

46U3.11,Urine culture - Escherich.coli

46U4.00,Urine culture - Proteus

46U5.00,Urine culture - Str. faecalis

46U6.00,Urine culture - Staph. albus

46U7.00,Urine culture - Pseudomonas

46U8.00,Urine culture - Bacteria OS

46X0.00,Urine nitrite positive

A994.00,Nonspecific urethritis

K10..00,Infections of kidney

K100.00,Chronic pyelonephritis

K100000,Chronic pyelonephritis without medullary necrosis

K100100,Chronic pyelonephritis with medullary necrosis

K100z00,Chronic pyelonephritis NOS

K101.00,Acute pyelonephritis

K101000,Acute pyelonephritis without medullary necrosis

K101100,Acute pyelonephritis with medullary necrosis

K101400,Emphysematous pyelonephritis

K101z00,Acute pyelonephritis NOS

K104.00,Xanthogranulomatous pyelonephritis

K10y.00,Pyelonephritis and pyonephrosis unspecified

K10y000,Pyelonephritis unspecified

K10y300,Pyelonephritis in diseases EC

K10yz00,Unspecified pyelonephritis NOS

K10z.00,Infection of kidney NOS

K15..00,Cystitis

K150.00,Acute cystitis

K151.00,Chronic interstitial cystitis

K151200,Submucous cystitis

K151z00,Chronic interstitial cystitis NOS

K152.00,Other chronic cystitis

K152000,Subacute cystitis

K152y00,Chronic cystitis unspecified

K152z00,Other chronic cystitis NOS

K153.11,Follicular cystitis

K154.00,Cystitis in diseases EC

K154z00,Cystitis in diseases EC NOS

K155.00,Recurrent cystitis

K15y.00,Other specified cystitis

K15y000,Cystitis cystica

K15y200,Abscess of bladder

K15yz00,Other cystitis NOS

K15z.00,Cystitis NOS

K17..00,Urethritis due to non venereal causes

K17..11,Periurethritis

K170.00,Urethral and periurethral abscess

K170.11,Urethral abscess

K170000,Urethral abscess unspecified

K170100,Bulbourethral gland abscess

K170200,Urethral gland abscess

K170311,Periurethritis

K170400,Periurethral abscess

K170z00,Urethral abscess NOS

K17y.00,Other urethritis

K17y000,Urethritis unspecified

K17yz00,Other urethritis NOS

K17z.00,Urethritis due to non venereal cause NOS

K190.00,"Urinary tract infection, site not specified"

K190.11,Recurrent urinary tract infection

K190000,"Bacteriuria, site not specified"

K190011,Asymptomatic bacteriuria

K190100,"Pyuria, site not specified"

K190300,Recurrent urinary tract infection

K190400,Chronic urinary tract infection

K190500,Urinary tract infection

K190z00,"Urinary tract infection, site not specified NOS"

Kyu5000,[X]Other chronic cystitis

Kyu5100,[X]Other cystitis

Kyu5500,[X]Other urethritis

Kyu5800,[X]Urethritis in diseases classified elsewhere

R081.00,[D]Dysuria

R081z00,[D]Dysuria NOS

Codes for gastrointestinal (GI) infections

code,frequency

A020.00,Salmonella gastroenteritis

A020.11,Salmonellosis

A0...13,Vomiting - infective

A00..00,Cholera

A00..11,Vibrio cholerae

A000.00,Cholera - Vibrio cholerae

A001.00,Cholera - Vibrio cholerae El Tor

A00z.00,Cholera NOS

A0...11,Bacterial food poisoning

A0...12,Food poisoning

A01..00,Typhoid and paratyphoid fevers

A010.00,Typhoid fever

A010.11,Enteric fever

A011.00,Paratyphoid fever A

A012.00,Paratyphoid fever B

A013.00,Paratyphoid fever C

A01z.00,Paratyphoid fever NOS

A02..00,Other salmonella infections

A020.12,Salmonella food poisoning

A021.00,Salmonella septicaemia

A022.00,Localised salmonella infection

A022000,Local salmonella infection unspecified

A02y.00,Other specified salmonella infection

A02z.00,Salmonella infection NOS

A03..00,Shigellosis

A030.00,Shigella dysenteriae (group A)

A030.11,Bacillary dysentery

A031.00,Shigella flexneri (group B)

A032.00,Shigella boydii (group C)

A033.00,Shigella sonnei (group D)

A033.11,Bacillary dysentery Shigella sonnei

A03y.00,Other specified shigella infection

A03z.00,Shigellosis NOS

A04..00,Other bacterial food poisoning

A040.00,Staphylococcal food poisoning

A042.00,Clostridium perfringens food poisoning

A043.00,Other clostridia causing food poisoning

A044.00,Vibrio parahaemolyticus food poisoning

A04y.00,Other specified bacterial food poisoning

A04y000,Foodborne Bacillus cereus intoxication

A04z.00,Food poisoning NOS

A05..00,Amoebiasis

A050.00,Acute amoebic dysentery

A051.00,Chronic intestinal amoebiasis

A05y200,Amoeboma

A05yz00,Amoebic infection of other sites NOS

A05z.00,Amoebiasis NOS

A06..00,Other protozoal intestinal diseases

A060.00,Balantidiasis

A061.00,Giardiasis - Lambliasis

A061.11,Colitis - giardial

A062.00,Coccidiosis

A063.00,Intestinal trichomoniasis

A064.00,Cryptosporidiosis

A06y.00,Other specified protozoal intestinal diseases

A06z.00,Protozoal intestinal diseases NOS

A07..00,Intestinal infection due to other organisms

A070.00,Escherichia coli gastrointestinal tract infection

A070000,Enteropathogenic Escherichia coli infection

A070100,Enterotoxigenic Escherichia coli infection

A070200,Enteroinvasive Escherichia coli infection

A070300,Enterohaemorrhagic Escherichia coli infection

A071.00,Arizona paracolon gastrointestinal tract infection

A072.00,Aerobacter aerogenes gastrointestinal tract infection

A073.00,Proteus gastrointestinal tract infection

A073000,Proteus mirabilis gastrointestinal tract infection

A073100,Proteus morganii gastrointestinal tract infection

A073z00,Proteus gastrointestinal tract infection NOS

A074.00,Other specified gastrointestinal tract bacterial infection

A074000,Staphylococcal gastrointestinal tract infection

A074011,Diarrhoea due to staphylococcus

A074012,Diarrhoea due to staphylococcal toxin

A074100,Pseudomonas gastrointestinal tract infection

A074111,Diarrhoea due to Pseudomonas pyocyanea

A074300,Campylobacter gastrointestinal tract infection

A074311,Diarrhoea due to Campylobacter jejuni

A074312,Campylobacter enteritis

A074313,Helicobacter gastritis

A074400,Enteritis due to Yersinia enterocolitica

A074500,Helicobacter pylori gastrointestinal tract infection

A074y00,Other specified other gastrointestinal infection

A074z00,Other specified gastrointestinal tract infections NOS

A075.00,Unspecified bacterial enteritis

A076.00,Enteritis due to specified virus

A076.11,Viral diarrhoea

A076.12,Viral vomiting

A076000,Enteritis due to adenovirus

A076100,Enteritis due to enterovirus

A076200,Enteritis due to rotavirus

A076300,Enteritis due to norovirus

A076z00,Enteritis due to specified virus NOS

A07y.00,Gastrointestinal tract infection specified organism NEC

A07y000,Viral gastroenteritis

A07y100,Infantile viral gastroenteritis

A07z.00,Gastrointestinal tract infection specified organism NOS

A08..00,Ill-defined intestinal tract infections

A08..11,Gastric flu

A080.00,"Infectious colitis, enteritis and gastroenteritis"

A080200,Infectious enteritis

A080300,Infectious gastroenteritis

A080z00,"Infectious colitis, enteritis and gastroenteritis NOS"

A081.00,"Colitis, enteritis and gastroenteritis presumed infectious"

A081.11,"Colitis,enteritis ? infectious"

A081000,Colitis - presumed infectious origin

A081100,Enteritis - presumed infectious origin

A081200,Gastroenteritis - presumed infectious origin

A081z00,"Colitis, enteritis and gastroenteritis presumed infect NOS"

A082.00,Infectious diarrhoea

A082.11,Travellers' diarrhoea

A082000,Dysenteric diarrhoea

A082100,Epidemic diarrhoea

A082111,Viral gastroenteritis

A082z00,Infectious diarrhoea NOS

A083.00,Diarrhoea of presumed infectious origin

A083.11,Diarrhoea & vomiting -? infect

A08z.00,Ill defined gastrointestinal tract infections NOS

A0y..00,Other specified infectious diseases of intestinal tract

A0z..00,Intestinal tract infectious disease NOS

Ayu0.00,[X]Intestinal infectious diseases

Ayu0000,"[X]Cholera, unspecified"

Ayu0100,"[X]Paratyphoid fever, unspecified"

Ayu0200,[X]Other specified salmonella infections

Ayu0300,"[X]Salmonella infection, unspecified"

Ayu0400,[X]Other shigellosis

Ayu0500,"[X]Shigellosis, unspecified"

Ayu0600,[X]Other specified bacterial intestinal infections

Ayu0700,"[X]Bacterial intestinal infection, unspecified"

Ayu0800,[X]Other specified bacterial food-borne intoxications

Ayu0900,"[X]Bacterial food-borne intoxication, unspecified"

Ayu0B00,"[X]Amoebiasis, unspecified"

Ayu0C00,[X]Other specified protozoal intestinal diseases

Ayu0D00,"[X]Protozoal intestinal disease, unspecified"

Ayu0E00,[X]Other viral enteritis

Ayu0F00,"[X]Viral intestinal infection, unspecified"

Ayu0G00,[X]Other specified intestinal infections

Ayu0H00,[X]Diarrhoea+gastroenteritis of presumed infectious origin

J43..11,Gastroenteritis

A0...00,Intestinal infectious diseases

A052.00,Amoebic nondysenteric colitis

A080100,Infectious colitis

A080400,Catarrhal dysentery

A080500,Haemorrhagic dysentery

A083.12,``

**Suppl. Table 1:** Absolute and relative risk of infections in AD patients with at least two prescriptions of mineralocorticoids and glucocorticoids and matched cohort.

|  | **AD cohort (n = 1493)** | **Matched unexposed cohort (n = 2984)** |
| --- | --- | --- |
| **Lower respiratory tract infections** | | |
| Outcome events, n. (%) | 124 (8.31) | 129 (4.32) |
| Person-years | 10189 | 22106 |
| Crude incidence rate/1000-person years | 12.17 | 5.83 |
| Follow-up years, median (IQR) | 5.18 (2.07-10.69) | 6.03 (2.55-11.31) |
| Unadjusted incidence rate ratio (95% CI)  p-value | 2.08 (1.63-2.67)  p <0.001 | |
| Adjusted incidence rate ratio (95% CI)^†^  p-value | 2.10 (1.63-2.71)  p <0.001 | |
| **Urinary tract infections** | | |
| Outcome events, n. (%) | 268 (17.95) | 380 (12.73) |
| Person-years | 9130 | 20301 |
| Crude incidence rate/1000-person years | 29.35 | 18.72 |
| Follow-up years, median (IQR) | 4.49 (1.69-9.52) | 5.19 (2.18-10.45) |
| Unadjusted incidence rate ratio (95% CI)  p-value | 1.57 (1.34-1.83)  p <0.001 | |
| Adjusted incidence rate ratio (95% CI)^†^  p-value | 1.46 (1.25-1.72)  p <0.001 | |
| **Gastro-intestinal infections** | | |
| Outcome events, n. (%) | 192 (1286) | 101 (3.38) |
| Person-years | 9455 | 21959 |
| Crude incidence rate/1000-person years | 20.03 | 4.60 |
| Follow-up years, median (IQR) | 4.73 (1.81-9.72) | 5.90 (2.55-11.20) |
| Unadjusted incidence rate ratio (95% CI)  p-value | 4.41 (3.47-5.62)  p <0.001 | |
| Adjusted incidence rate ratio (95% CI)^†^  p-value | 4.07 (3.17-5.21)  p <0.001 | |

^†^Adjusted for age, gender, smoking status, BMI, Townsend Deprivation Index and Charlson Comorbidity Index.

**Suppl. Table 2**: Antimicrobial prescriptions counts in AD patients with at least two prescriptions of mineralocorticoids and glucocorticoids compared to the matched control cohort.

|  | **AD cohort (n = 1493)** | **Matched unexposed cohort (n = 2984)** |
| --- | --- | --- |
| **Antibiotic prescriptions** | | |
| Count of prescriptions, n. | 12819 | 15408 |
| Person-years | 10609 | 22554 |
| Count rates (per 1000 years) | 1208 | 683 |
| Follow-up years, median (IQR) | 5.50 (2.21-11.07) | 6.12 (2.67-6.12) |
| Unadjusted incidence rate ratio (95% CI)  p-value | 1.77 (1.73-1.81)  p <0.001 | |
| Adjusted incidence rate ratio (95% CI)^†^  p-value | 1.68 (1.64-1.73)  p <0.001 | |
| **Antifungal prescriptions** | | |
| Count of prescriptions, n. | 1157 | 1185 |
| Person-years | 10609 | 22554 |
| Count rates (per 1000 years) | 109 | 52 |
| Follow-up years, median (IQR) | 5.50 (2.21-11.07) | 6.12 (2.67-6.12) |
| Unadjusted incidence rate ratio (95% CI)  p-value | 2.08 (1.91-2.25)  p <0.001 | |
| Adjusted incidence rate ratio (95% CI)^†^  p-value | 1.83 (1.68-1.99)  p <0.001 | |

^†^Adjusted for age, gender, smoking status, BMI, Townsend Deprivation Index and Charlson Comorbidity Index.

**Suppl. Table 3:** Absolute and relative risk of infections in AD patients with and without type 1 diabetes mellitus and matched controls.

|  | **Patients with type 1 diabetes mellitus** | | **Patients without type 1 diabetes mellitus** | |
| --- | --- | --- | --- | --- |
|  | **AD cohort (n = 127)** | **Matched unexposed**  **cohort (n = 254)** | **AD cohort (n = 1453)** | **Matched unexposed**  **cohort (n = 2904)** |
| **Lower respiratory tract infections** | | |  |  |
| Outcome events, n. (%) | 13 (10.24) | 11 (4.33) | 117 (8.05) | 126 (4.34) |
| Person-years | 762 | 1789 | 9575 | 21047 |
| Crude incidence rate/1000-person years | 17.06 | 6.15 | 12.22 | 5.99 |
| Follow-up years, median (IQR) | 5.11 (1.40-9.40) | 6.06 (1.80-10.75) | 4.85 (1.79-10.31) | 5.75 (2.42-11.15) |
| Unadjusted incidence rate ratio (95% CI)  p-value | 2.77 (1.24-6.19)  p =0.013 | | 2.04 (1.59-2.62)  p <0.001 | |
| Adjusted incidence rate ratio (95% CI)^†^  p-value | 2.24 (0.64-7.89)  P=0.208 | | 2.11 (1.63-2.73)  p <0.001 | |
| **Urinary tract infections** | | |  |  |
| Outcome events, n. (%) | 22 (17.32) | 26 (10.24) | 260 (17.89) | 370 (12.74) |
| Person-years | 676 | 1662 | 8572 | 19341 |
| Crude incidence rate/1000-person years | 32.56 | 15.64 | 30.33 | 19.13 |
| Follow-up years, median (IQR) | 3.64 (0.97-8.71) | 4.95 (1.50-10.10) | 4.11 (1.50-9.25) | 5.08 (2.13-10.27) |
| Unadjusted incidence rate ratio (95% CI)  p-value | 2.08 (1.18-3.67)  p =0.011 | | 1.58 (1.35-1.86)  p <0.001 | |
| Adjusted incidence rate ratio (95% CI)^†^  p-value | 1.27 (0.57-2.85)  p =0.556 | | 1.52 (1.29-1.79)  p <0.001 | |
| **Gastro-intestinal infections** | | |  |  |
| Outcome events, n. (%) | 23 (18.11) | 10 (3.94) | 171 (11.77) | 100 (3.44) |
| Person-years | 677 | 1789 | 8920 | 20875 |
| Crude incidence rate/1000-person years | 33.97 | 5.60 | 19.17 | 4.79 |
| Follow-up years, median (IQR) | 3.42 (1.08-7.90) | 5.69 (1.62-10.75) | 4.52 (1.70-9.40) | 5.62 (2.44-11.05) |
| Unadjusted incidence rate ratio (95% CI)  p-value | 6.07 (2.89-12.75)  p <0.001 | | 4.00 (3.13-5.12)  p <0.001 | |
| Adjusted incidence rate ratio (95% CI)^†^  p-value | 6.16 (1.85-20.51)  p =0.003 | | 3.74 (2.91-4.82)  p <0.001 | |

^†^Adjusted for age, gender, smoking status, BMI, Townsend Deprivation Index and Charlson Comorbidity Index.

**Suppl. Table 4**: Antimicrobial prescriptions counts in AD patients with and without type 1 diabetes mellitus and matched controls.

|  | **Patients with Type 1 diabetes mellitus** | | **Patients without Type 1 diabetes mellitus** | |
| --- | --- | --- | --- | --- |
|  | **AD cohort (n = 127)** | **Matched unexposed**  **cohort (n = 254)** | **AD cohort (n = 1453)** | **Matched unexposed**  **cohort (n = 2904)** |
| **Antibiotic prescriptions** | | |  |  |
| Count of prescriptions, n. | 1162 | 1101 | 12124 | 14789 |
| Person-years | 818 | 1840 | 9949 | 21467 |
| Count rates (per 1000 years) | 1420 | 598 | 1219 | 689 |
| Follow-up years, median (IQR) | 5.16 (1.40-10.32) | 6.09 (1.80-11.12) | 5.11 (1.99-10.79) | 5.88 (2.62-11.34) |
| Unadjusted incidence rate ratio (95% CI)  p-value | 2.37 (2.19-2.58)  p<0.001 | | 1.77 (1.73-1.81)  p <0.001 | |
| Adjusted incidence rate ratio (95% CI)^†^  p-value | 1.28 (1.23-1.54)  P<0.001 | | 1.73 (1.69-1.77)  p <0.001 | |
| **Antifungal prescriptions** | | |  |  |
| Count of prescriptions, n. | 97 | 126 | 1094 | 1087 |
| Person-years | 818 | 1840 | 9949 | 21467 |
| Count rates (per 1000 years) | 119 | 68 | 110 | 51 |
| Follow-up years, median (IQR) | 5.16 (1.40-10.32) | 6.09 (1.80-11.12) | 5.11 (1.99-10.79) | 5.88 (2.62-11.34) |
| Unadjusted incidence rate ratio (95% CI)  p-value | 1.73 (1.33-2.26)  p <0.001 | | 2.17 (2.00-2.36)  p <0.001 | |
| Adjusted incidence rate ratio (95% CI)^†^  p-value | 2.28 (1.49-3.49)  p <0.001 | | 2.02 (1.85-2.20)  p <0.001 | |

^†^Adjusted for age, gender, smoking status, BMI, Townsend Deprivation Index and Charlson Comorbidity Index.
